# Gliotoxin, a known virulence factor in the major human pathogen *Aspergillus fumigatus*, is also biosynthesized by the non-pathogenic relative *A. fischeri*

**DOI:** 10.1101/868745

**Authors:** Sonja L. Knowles, Matthew E. Mead, Lilian Pereira Silva, Huzefa A. Raja, Jacob L. Steenwyk, Gustavo H. Goldman, Antonis Rokas, Nicholas H. Oberlies

**Affiliations:** Department of Chemistry and Biochemistry, University of North Carolina at Greensboro, Greensboro, North Carolina, USA; Department of Biological Sciences, Vanderbilt University, Nashville, Tennessee, USA; Faculdade de Ciencias Farmacêuticas de Ribeirão Preto, Universidade de São Paulo, Brazil

**Author notes:** **CORRESPONDING AUTHORS:** Antonis Rokas, Gustavo H. Goldman, and Nicholas H. Oberlies.

**Keywords:** fungal pathogenesis, secondary metabolism, gliotoxin, specialized metabolism, evolution of virulence, *laeA*, aspergillosis

## Abstract

*Aspergillus fumigatus* is a major opportunistic human pathogen. Multiple traits contribute to *A. fumigatus* pathogenicity, including its ability to produce specific secondary metabolites, such as gliotoxin. Gliotoxin is known to inhibit the host immune response, and genetic mutants that inactivate gliotoxin biosynthesis (or secondary metabolism in general) attenuate *A. fumigatus* virulence. The genome of *A. fischeri*, a very close non-pathogenic relative of *A. fumigatus*, contains a biosynthetic gene cluster that exhibits high sequence similarity to the *A. fumigatus* gliotoxin cluster. However, *A. fischeri* is not known to produce gliotoxin. To gain further insight into the similarities and differences between the major pathogen *A. fumigatus* and the non-pathogen *A. fischeri*, we examined whether *A. fischeri* strain NRRL 181 biosynthesizes gliotoxin and whether its production, and of secondary metabolites more generally, influence its virulence profile. We found that *A. fischeri* biosynthesizes gliotoxin in the same conditions as *A. fumigatus*. However, whereas loss of *laeA*, a master regulator of secondary metabolite production, has been previously shown to reduce the virulence of *A. fumigatus*, we found that *laeA* loss (and loss of secondary metabolite production, including gliotoxin) in *A. fischeri* does not influence its virulence. These results suggest that gliotoxin and secondary metabolite production are virulence factors in the genomic and phenotypic background of the major pathogen *A. fumigatus* but are much less important in the background of the non-pathogen *A. fischeri*. We submit that understanding the observed spectrum of pathogenicity across closely related pathogenic and non-pathogenic *Aspergillus* species will require detailed characterization of their biological, chemical, and genomic similarities and differences.

**IMPORTANCE:** *Aspergillus fumigatus* is a major opportunistic fungal pathogen of humans but most of its close relatives are non-pathogenic. Why is that so? This important, yet largely unanswered, question can be addressed by examining how *A. fumigatus* and its non-pathogenic close relatives are similar or different with respect to virulence-associated traits. We investigated whether *Aspergillus fischeri*, a non-pathogenic close relative of *A. fumigatus*, can produce gliotoxin, a mycotoxin known to contribute to *A. fumigatus* virulence. We discovered that the non-pathogenic *A. fischeri* produces gliotoxin under the same conditions as the major pathogen *A. fumigatus*. However, we also discovered that, in contrast to what has been previously observed in *A. fumigatus*, loss of secondary metabolite, including gliotoxin, production in *A. fischeri* does not alter its virulence. Our results are consistent with the “cards of virulence” model of opportunistic fungal disease, where the ability to cause disease stems from the combination (“hand”) of individual virulence factors (“cards”), but not from individual factors *per se*.

---

*Aspergillus fumigatus* is a major fungal pathogen responsible for hundreds of thousands of infections and deaths each year (1, 2). Several secondary metabolites biosynthesized by *A. fumigatus* have been shown to be required for disease (3). For example, gliotoxin (**Fig. 1A**), a secondary metabolite that belongs to the epipolythiodioxopiperazine (ETP) class of mycotoxins (4), can be detected in the sera of patients with invasive aspergillosis (5) and is known to inhibit the host immune response (3). When the *gliP* gene, which encodes the major synthetase of the gliotoxin biosynthetic gene cluster, is deleted from *A. fumigatus*, the mutant strain does not biosynthesize gliotoxin and exhibits attenuated virulence in a non-neutropenic murine model of aspergillosis (6–8). Similarly, deletion of *laeA*, a positive regulator of several *A. fumigatus* secondary metabolites, including gliotoxin, also reduces virulence (9, 10). These results establish that gliotoxin, as well as other secondary metabolites, contribute to *A. fumigatus* virulence (3).

**FIG 1.**
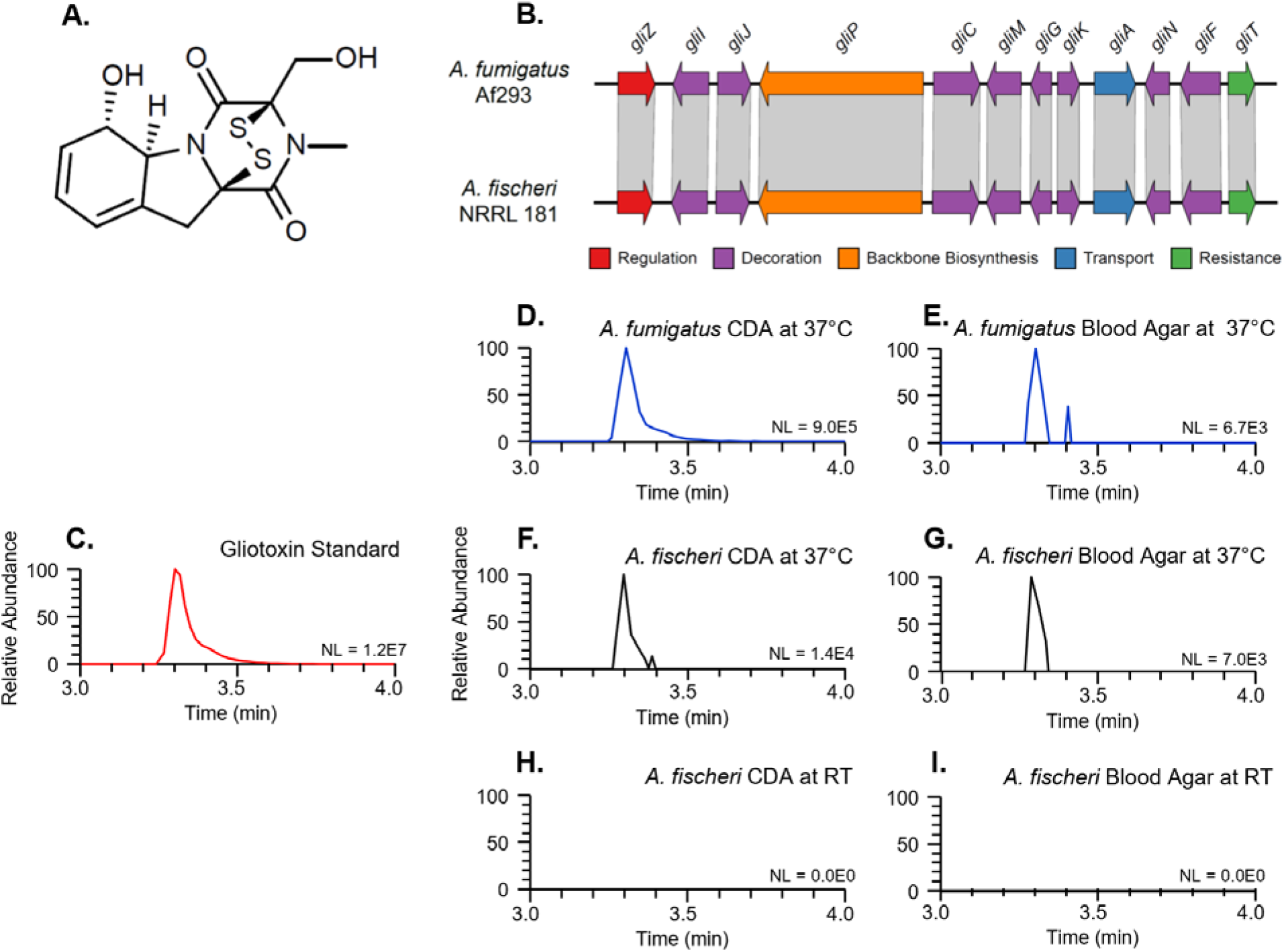
*Aspergillus fischeri* biosynthesizes gliotoxin when grown in conditions that induce *A. fumigatus* gliotoxin biosynthesis. **A.** Chemical structure of gliotoxin. **B.** The genome of the non-pathogenic species *A. fischeri* strain NRRL 181 (12, 19) contains a biosynthetic gene cluster homologous to the gliotoxin cluster in the major pathogen *A. fumigatus* strain Af293 (4). Arrows indicate genes and the direction in which they are transcribed. Red – regulation, Purple – decoration, Orange – backbone biosynthesis, Blue – transport, Green – resistance. Homologous genes are connected by gray parallelograms. **C – I.** Chromatograms demonstrating the biosynthesis of gliotoxin in *A. fischeri* when grown on Czapek-Dox Agar (CDA) or blood agar at 37°C. Each sample (0.2 mg/mL) was analyzed by UHPLC-HRESIMS, and the data are presented as extracted ion chromatograms (XIC) using the protonated mass of gliotoxin (C_13_H_15_N_2_O_4_S_2_; [M+H]^+^ = 327.0473) and a window of ± 5.0 ppm. **C.** Analysis of the gliotoxin standard (0.01 mg/mL). **D.** *A. fumigatus* grown on CDA at 37°C. **E.** *A. fumigatus* grown on blood agar at 37°C. **F.** *A. fischeri* grown on CDA incubated at 37°C. **G.** *A. fischeri* grown on blood agar incubated at 37°C. **H.** *A. fischeri* grown on CDA at room temperature (RT). **I.** *A. fischeri* grown on blood agar at RT. The retention time (3.30 min) and accurate mass (327.0473 ± 5.0 ppm) data confirm the biosynthesis of gliotoxin by *A. fischeri* in panels F and G. NL: Normalization Level (i.e., base peak intensity; the larger the NL value the better the signal to noise ratio).

Even though *A. fumigatus* is a major pathogen, its closest relatives are non-pathogenic (11–13). For example, the closely related species *Aspergillus fischeri* has been identified as the cause of only a handful of clinical cases (14, 15) and is not considered pathogenic. Detailed comparisons of virulence in diverse murine and invertebrate models of fungal disease have shown that *A. fischeri* is much less virulent than *A. fumigatus* (12). It is important to emphasize here that in diverse animal models of aspergillosis, especially when high inoculums of spores are administered, non-pathogens can sometimes cause disease, as we have observed in previous experiments with *A. fischeri*; however, in all such cases non-pathogens exhibit lower levels of virulence than pathogens (12).

Despite their significant differences in ability to cause fungal disease, a recent examination of known genetic contributors to virulence revealed that nearly all genes known to contribute to *A. fumigatus* disease are also present in *A. fischeri* (12). For example, both species appear to contain *laeA*, and deletion of the *laeA* gene in either species is known to reduce biosynthesis of secondary metabolites (12, 16), suggesting that the gene’s function is conserved. Similarly, both species appear to contain intact gliotoxin biosynthetic gene clusters (**Fig. 1B**); however, gliotoxin production has been shown in *A. fumigatus* and a few other closely related species (17), but not in *A. fischeri* (12, 17). These data raise two questions: Is *A. fischeri* capable of biosynthesizing gliotoxin? And if it is, how does production of gliotoxin, and secondary metabolites more generally, influence its virulence profile?

## *A. fischeri*, a non-pathogenic relative of the major pathogen *A. fumigatus*, can also biosynthesize gliotoxin

To test whether *A. fischeri* biosynthesizes gliotoxin, we examined the chemical profile of a standard of gliotoxin and extracts of *A. fumigatus* strain Af293, and *A. fischeri* strain NRRL 181 via UHPLC–HRESIMS (ultra-high-performance liquid chromatography–high-resolution electrospray ionization mass spectrometry) (**Supplementary Methods**). We collected three sets of data, specifically chromatographic retention time, high resolution mass spectrometry data, and tandem mass spectrometry fragmentation patterns. Analysis of a gliotoxin standard (**Fig. 1C**) showed that it elutes at 3.30 minutes, with an accurate mass of 327.0464 Da (2.8 ppm) and has key fragments of 263.1 Da and 245.1 Da, in accord with values reported in the literature (18).

We next analyzed *A. fumigatus* strain Af293, which we used as a positive control, since it is known to biosynthesize gliotoxin (17). When *A. fumigatus* was grown on Czapek-Dox Agar (CDA) at 37°C (**Fig. 1D**), a peak with the same retention time (3.30 min), HRESIMS spectrum, MS/MS spectrum, and accurate mass of 327.0463 (3. 1ppm) was noted (**Figs. S1 and S2**). We also detected gliotoxin production, albeit in lower abundance, when we cultured *A. fumigatus* on 5% blood agar at 37°C (**Fig. 1E**). In contrast, we did not observe gliotoxin production when we grew *A. fumigatus* on oatmeal agar at 37°C (**Fig. S3**).

To test whether *A. fischeri* biosynthesized gliotoxin, we grew strain NRRL 181 on the same media and temperature conditions as *A. fumigatus*. When *A. fischeri* was grown on CDA at 37°C, we observed a peak with the same retention time (**Fig. 1F**), HRESIMS spectrum (**Fig. S4**), and MS/MS spectrum as that of *A. fumigatus* (**Fig. S5**), indicating gliotoxin biosynthesis in *A. fischeri*. Similarly, we detected gliotoxin production in lower abundance when we grew *A. fischeri* on 5% blood agar at 37°C (**Fig. 1G**). In contrast, we did not observe gliotoxin production when we grew *A. fischeri* on CDA or on 5% blood agar at room temperature (**Figs. 1H** and **1I**, respectively) or on oatmeal agar at 37°C (**Fig. S3**). These results demonstrate that: a) the non-pathogen *A. fischeri* biosynthesizes similar quantities of gliotoxin in the same conditions that induce gliotoxin biosynthesis in the major pathogen *A. fumigatus*, and b) similar to what has been previously observed in *A. fumigatus* (20), both growth medium and temperature influence gliotoxin biosynthesis in *A. fischeri*.

## *laeA*, a master regulator of secondary metabolism and *A. fumigatus* virulence factor, is not a virulence factor in *A. fischeri*

To test whether the regulation of gliotoxin production, and more generally of secondary metabolite production, contributes to the virulence profile of *A. fischeri*, we deleted the endogenous copy of *laeA* from *A. fischeri* and infected larvae of the moth *Galleria mellonella*, a well-established invertebrate model of fungal disease (21), with the resulting mutant strain (Supplementary Methods). Infection of *G. mellonella* larvae with *A. fumigatus* is known to induce gliotoxin biosynthesis (22). We infected asexual spores (conidia) at two different concentrations and compared the survival curves between the Δ*laeA* and the wild-type (WT) strain of *A. fischeri* (Fig. 2). At both concentrations, our experiments showed that moth larval survival was not significantly different between the Δ*laeA* and the WT strains.

**FIG 2.**
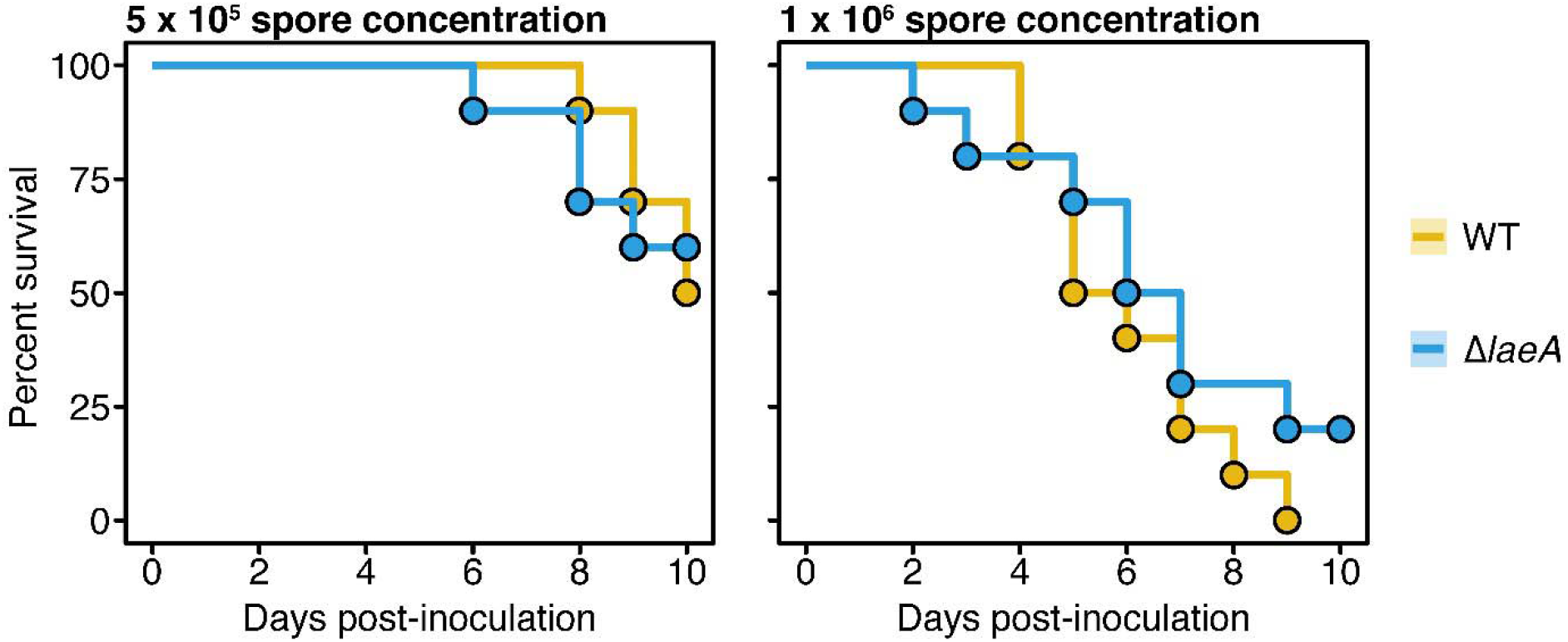
Deletion of the master regulator *laeA* in *A. fischeri* does not alter its virulence. Cumulative survival of moth *(Galleria mellonella)* larvae inoculated with 5 × 10^5^ (left) or 1 × 10^6^ (right) asexual spores or conidia of either the Δ*laeA* mutant or the wild-type (WT) *A. fischeri* NRRL 181 strain. Comparisons of moth cumulative survival when infected with either Δ*laeA* or WT strain revealed no statistically significant differences at spore concentrations of 5 × 10^5^ (left) or 1 × 10^6^ (right) (*p*-value = 0.91 and 0.30, respectively; log-rank test). For the inoculations, ten moths were infected per group.

Importantly, the Δ*laeA* strain of *A. fischeri* NRRL 181 is known to exhibit reduced production of secondary metabolites in diverse conditions in a manner consistent with the gene’s role as a master regulator of secondary metabolism (12). To confirm that the Δ*laeA* strain does not produce gliotoxin, we analyzed it using the same chemical methods that showed production of gliotoxin in the WT strain following growth on CDA or 5% blood agar at 37°C. In contrast to the WT strain (Figs. **1F** and **1G**), we did not observe gliotoxin production in the Δ*laeA* strain (**Fig. S3**). Whereas loss of *laeA* and secondary metabolite – including gliotoxin – production has been previously shown to reduce the virulence of the major pathogen *A. fumigatus* (9, 10), our results suggest that loss of *laeA* (and secondary metabolite production, including gliotoxin) in *A. fischeri* does not influence its virulence.

## Concluding remarks

In this study, we show for the first time that *A. fischeri –* when grown in conditions known to induce gliotoxin production in *A. fumigatus* – can biosynthesize gliotoxin (**Fig. 1**). Furthermore, we show that an *A. fischeri* mutant that lacks a master regulatory gene of secondary metabolism (*laeA*) does not alter the pathogenic potential of *A. fischeri* (**Fig. 2**). Thus, it appears that gliotoxin and secondary metabolite production more generally are virulence factors in the genomic and phenotypic background of the pathogen *A. fumigatus* but that they are much less important for virulence in the genomic background of the non-pathogen *A. fischeri*. These results provide support for the “cards of virulence” model of opportunistic fungal disease (23), where the ability to cause disease stems from the combination (“hand”) of individual virulence factors (“cards”). We hypothesize that while *A. fischeri* possesses the “cards” for gliotoxin production and secondary metabolism regulation, its cumulative “hand” is thankfully not a winner when it comes to causing disease.

## SUPPLEMENTAL MATERIAL

Single supplementary document that contains additional experimental methods and five supplementary figures. The figures are:

**FIG S1.** The mass spectra of gliotoxin in *A. fumigatus* grown on CDA and blood agar at 37°C verify the biosynthesis of gliotoxin by cultures of *A. fumigatus* on both CDA and blood agar at 37°C. Data are presented as mass to charge ratios (*m/z*).

**FIG S2.** The fragmentation pattern (i.e., MS/MS data) of gliotoxin in *A. fumigatus* grown on CDA and blood agar at 37°C verify the biosynthesis of gliotoxin in both the CDA and blood agar growths of *A. fumigatus* at 37°C.

**FIG S3.** Base peak chromatograms of the gliotoxin standard, *A. fumigatus* grown on OMA at 37°C, *A. fischeri* grown on OMA at 37°C, Δ*laeA A. fischeri* grown on CDA at 37°C, and Δ*laeA A. fischeri* grown on blood agar at 37°C show that some media do not induce gliotoxin biosynthesis.

**FIG S4.** The mass spectra of gliotoxin in *A. fischeri* verify the biosynthesis of gliotoxin by cultures of *A. fischeri* on both CDA and blood agar at 37°C.

**FIG S5.** The fragmentation patterns (i.e., MS/MS data) of gliotoxin in *A. fischeri* further verify the biosynthesis of gliotoxin by cultures of *A. fischeri* in both the CDA and blood agar at 37°C.

## ACKNOWLEDGEMENTS

Computational infrastructure was provided by The Advanced Computing Center for Research and Education (ACCRE) at Vanderbilt University. S.L.K. was supported by the National Center for Complementary and Integrative Health, a component of the National Institutes of Health, under award number F31 AT010558. M.E.M. and A.R. were supported by a Vanderbilt University Discovery Grant; J.L.S. and A.R. were supported by the Howard Hughes Medical Institute through the James H. Gilliam Fellowships for Advanced Study program. Research in A.R.’s lab is also supported by the National Science Foundation (DEB-1442113), the Guggenheim Foundation, and the Burroughs Wellcome Fund. G.H.G. and L.P.S. were supported by grants from Fundação de Amparo à Pesquisa do Estado de São Paulo (FAPESP 2016/07870-9 and 2016/21392-2) and Conselho Nacional de Desenvolvimento Científico e Tecnológico (CNPq), both from Brazil.

## References

1. Bongomin F, Gago S, Oladele RO, Denning DW. 2017. Global and multi-national prevalence of fungal diseases-estimate precision. J Fungi (Basel) 3:57.

2. Brown GD, Denning DW, Gow NA, Levitz SM, Netea MG, White TC. 2012. Hidden killers: human fungal infections. Sci Transl Med 4:165rv13.

3. Raffa N, Keller NP. 2019. A call to arms: Mustering secondary metabolites for success and survival of an opportunistic pathogen. PLoS Pathog 15:e1007606.

4. Gardiner DM, Howlett BJ. 2005. Bioinformatic and expression analysis of the putative gliotoxin biosynthetic gene cluster of *Aspergillus fumigatus*. FEMS Microbiol Lett 248:241–8.

5. Lewis RE, Wiederhold NP, Chi J, Han XY, Komanduri KV, Kontoyiannis DP, Prince RA. 2005. Detection of gliotoxin in experimental and human aspergillosis. Infect Immun 73:635–7.

6. Sugui JA, Pardo J, Chang YC, Zarember KA, Nardone G, Galvez EM, Mullbacher A, Gallin JI, Simon MM, Kwon-Chung KJ. 2007. Gliotoxin is a virulence factor of *Aspergillus fumigatus: gliP* deletion attenuates virulence in mice immunosuppressed with hydrocortisone. Eukaryot Cell 6:1562–9.

7. Spikes S, Xu R, Nguyen CK, Chamilos G, Kontoyiannis DP, Jacobson RH, Ejzykowicz DE, Chiang LY, Filler SG, May GS. 2008. Gliotoxin production in *Aspergillus fumigatus* contributes to host-specific differences in virulence. The Journal of Infectious Diseases 197:479–86.

8. Cramer RA, Jr., Gamcsik MP, Brooking RM, Najvar LK, Kirkpatrick WR, Patterson TF, Balibar CJ, Graybill JR, Perfect JR, Abraham SN, Steinbach WJ. 2006. Disruption of a nonribosomal peptide synthetase in *Aspergillus fumigatus* eliminates gliotoxin production. Eukaryot Cell 5:972–80.

9. Bok JW, Balajee SA, Marr KA, Andes D, Nielsen KF, Frisvad JC, Keller NP. 2005. LaeA, a regulator of morphogenetic fungal virulence factors. Eukaryot Cell 4:1574–82.

10. Perrin RM, Fedorova ND, Bok JW, Cramer RA, Wortman JR, Kim HS, Nierman WC, Keller NP. 2007. Transcriptional regulation of chemical diversity in *Aspergillus fumigatus* by LaeA. PLoS Pathog 3:e50.

11. Sugui JA, Peterson SW, Figat A, Hansen B, Samson RA, Mellado E, Cuenca-Estrella M, Kwon-Chung KJ. 2014. Genetic relatedness versus biological compatibility between Aspergillus fumigatus and related species. J Clin Microbiol 52:3707–21.

12. Mead ME, Knowles SL, Raja HA, Beattie SR, Kowalski CH, Steenwyk JL, Silva LP, Chiaratto J, Ries LNA, Goldman GH, Cramer RA, Oberlies NH, Rokas A. 2019. Characterizing the pathogenic, genomic, and chemical traits of *Aspergillus fischeri*, a close relative of the major human fungal pathogen *Aspergillus fumigatus*. mSphere 4:e00018–19.

13. Hubka V, Barrs V, Dudova Z, Sklenar F, Kubatova A, Matsuzawa T, Yaguchi T, Horie Y, Novakova A, Frisvad JC, Talbot JJ, Kolarik M. 2018. Unravelling species boundaries in the *Aspergillus viridinutans* complex (section *Fumigati):* opportunistic human and animal pathogens capable of interspecific hybridization. Persoonia 41:142–174.

14. Lonial S, Williams L, Carrum G, Ostrowski M, McCarthy P, Jr. 1997. *Neosartorya fischeri:* an invasive fungal pathogen in an allogeneic bone marrow transplant patient. Bone Marrow Transplant 19:753–5.

15. Coriglione G, Stella G, Gafa L, Spata G, Oliveri S, Padhye AA, Ajello L. 1990. *Neosartorya fischeri* var *fischeri* (Wehmer) Malloch and Cain 1972 (anamorph: *Aspergillus fischerianus* Samson and Gams 1985) as a cause of mycotic keratitis. Eur J Epidemiol 6:382–5.

16. Bok JW, Keller NP. 2004. LaeA, a regulator of secondary metabolism in *Aspergillus* spp. Eukaryot Cell 3:527–35.

17. Frisvad JC, Larsen TO. 2015. Extrolites of *Aspergillus fumigatus* and other pathogenic species in *Aspergillus* section *Fumigati*. Front Microbiol 6:1485.

18. Pena GA, Monge MP, Gonzalez Pereyra ML, Dalcero AM, Rosa CA, Chiacchiera SM, Cavaglieri LR. 2015. Gliotoxin production by *Aspergillus fumigatus* strains from animal environment. Micro-analytical sample treatment combined with a LC-MS/MS method for gliotoxin determination. Mycotoxin Res 31:145–50.

19. Fedorova ND, Khaldi N, Joardar VS, Maiti R, Amedeo P, Anderson MJ, Crabtree J, Silva JC, Badger JH, Albarraq A, Angiuoli S, Bussey H, Bowyer P, Cotty PJ, Dyer PS, Egan A, Galens K, Fraser-Liggett CM, Haas BJ, Inman JM, Kent R, Lemieux S, Malavazi I, Orvis J, Roemer T, Ronning CM, Sundaram JP, Sutton G, Turner G, Venter JC, White OR, Whitty BR, Youngman P, Wolfe KH, Goldman GH, Wortman JR, Jiang B, Denning DW, Nierman WC. 2008. Genomic islands in the pathogenic filamentous fungus *Aspergillus fumigatus*. PLoS Genet 4:e1000046.

20. Belkacemi L, Barton RC, Hopwood V, Evans EG. 1999. Determination of optimum growth conditions for gliotoxin production by *Aspergillus fumigatus* and development of a novel method for gliotoxin detection. Med Mycol 37:227–33.

21. Fuchs BB, O’Brien E, El Khoury JB, Mylonakis E. 2010. Methods for using *Galleria mellonella* as a model host to study fungal pathogenesis. Virulence 1:475–482.

22. Reeves EP, Messina CG, Doyle S, Kavanagh K. 2004. Correlation between gliotoxin production and virulence of *Aspergillus fumigatus* in *Galleria mellonella*. Mycopathologia 158:73–9.

23. Casadevall A. 2006. Cards of virulence and the global virulome for humans. Microbe 1:359–364.

