## Supplementary Methods and Figures for "Gliotoxin, a known virulence factor in the major human pathogen *Aspergillus fumigatus*, is also biosynthesized by the non-pathogenic relative *A. fischeri*"

**SUPPLENTARY MATERIAL**

**Supplementary Methods**

**Fungal Strains.**	*Aspergillus fischeri* strain NRRL 181 was obtained from ARS Culture Collection (NRRL) (1). *A. fumigatus* strain Af293 was also utilized as a positive control (2).

**Growth Conditions.**	All strains were maintained on potato dextrose agar (PDA; Difco). To establish individual cultures, an agar square along with fungal mycelium was cut out aseptically from the leading edge of the culture and transferred onto blood agar (tryptic soy agar with 5% sheep’s blood; Hardy Diagnostics), Czapek-Dox agar (CDA; Difco), or oatmeal agar (OMA; Difco). All cultures at 37°C were maintained in an incubator (VWR International) in the dark over four days. All cultures at room temperature (RT; ~22°C) were kept for two weeks under 12h light/dark cycles. *A. fischeri* was grown on CDA (RT and 37°C), blood agar (RT and 37°C), and OMA (37°C). *A. fumigatus* was grown on CDA (37°C), blood agar (37°C), and OMA (37°C).

**Extraction.**	To evaluate the biosynthesis of gliotoxin in these fungal strains, cultures were extracted with organic solvents and analyzed by mass spectrometry (see below). The agar plates were extracted by spraying the fungal mycelium with MeOH, chopping it with a spatula, and transferring the contents to a scintillation vial. Acetone (~15 ml) was then added to the scintillation vial, and the resulting slurry was vortexed vigorously for approximately 3 min before steeping for 4 h at RT. Subsequently, the mixture was filtered and the resulting extract was dried under a stream of nitrogen gas.

**UHPLC-HRESIMS analysis.**	High-resolution electrospray ionization mass spectrometry (HRESIMS) experiments utilized a Thermo LTQ Orbitrap XL mass spectrometer (Thermo Fisher Scientific), equipped with an electrospray ionization source. This was coupled to an Acquity ultra-high-performance liquid chromatography (UHPLC) system (Waters Corp.), using a flow rate of 0.3 ml/min and a BEH C_18_ column (2.1 mm x 50 mm, 1.7 μm) that was operated at 40°C. The mobile phase consisted of CH_3_CN–H_2_O (Fischer Optima LC-MS grade; both acidified with 0.1% formic acid). The gradient began at 15% CH_3_CN and increased linearly to 100% CH_3­_CN over 8 mins, where it was held for 1.5 mins before returning to starting conditions to re-equilibrate.

Extracts were analyzed in the positive ion mode, scanning over a mass range of *m/z* 100 to 2,000 at a resolving power of 30,000. The spray voltage, source capillary, and tube lens voltages were set to 4.0 kV, 20 V, and 100 V, respectively, with a nitrogen sheath gas set to 30 arb and capillary temperature at 300°C. The fragmentation patterns (i.e., MS/MS data) were obtained by using an inclusion list containing the mass of gliotoxin ([M+H]^+^ = 327.047 *m/z*), with an isolation window of 2 Da and collision energy of 35%. The extracts and gliotoxin standard (Cayman Chemical Company) were prepared at a concentration of 0.2 mg/ml and 0.01 mg/ml, respectfully; both were dissolved in MeOH with an injection volume of 3 μl. To eliminate the possibility for sample carryover, two blanks (MeOH) were injected between every sample injection and the gliotoxin standard was analyzed at the end of the run.

**Virulence studies using an invertebrate model of fungal disease (*Galleria mellonella*).** These experiments were performed as previously described (1). Briefly, larvae of the moth *G. mellonella* were obtained by breeding adult moths (3) and selecting larvae that were similar in size (~275–330 mg). Prior to use, all larvae were kept for 24 hours in glass petri dishes in darkness at 37°C. Asexual spores (conidia) of Δ*laeA* mutant or the wild-type (WT) *A. fischeri* were obtained by growing the organism on a yeast extract-agar-glucose (YAG) medium for 2 days. Conidia were harvested in PBS and filtered through Miracloth (Calbiochem). Conidial concentration was estimated using a hemocytometer and conidial viability was assessed through incubation on YAG medium at 37°C for 48 h.

For infection assays, ten *G. mellonella* larvae in the final (sixth) instar larval stage of development were used per condition. Each larva in the test group was infected with a 5 μl inoculum of conidia from the Δ*laeA* mutant of *A. fischeri* (at either a 5 x 10^5^ spores / μl or a 1 x 10^6^ spores / μl concentration), whereas each larva in the control group was inoculated with the same concentration of the WT strain of *A. fischeri*. All inoculations were done using a Hamilton syringe (7000.5KH). All injections were performed at the hemocoel of each larva via the last left proleg. Following inoculation, all larvae were incubated in glass petri dishes in darkness at 37°C. Larval killing was scored daily. Larvae were considered dead by if they did not move in response to touch.

**Supplementary References**

1.	Mead ME, Knowles SL, Raja HA, Beattie SR, Kowalski CH, Steenwyk JL, Silva LP, Chiaratto J, Ries LNA, Goldman GH, Cramer RA, Oberlies NH, Rokas A. 2019. Characterizing the pathogenic, genomic, and chemical traits of *Aspergillus fischeri*, a close relative of the major human fungal pathogen *Aspergillus fumigatus*. mSphere 4:e00018-19.

2.	Nierman WC, Pain A, Anderson MJ, Wortman JR, Kim HS, Arroyo J, Berriman M, Abe K, Archer DB, Bermejo C, Bennett J, Bowyer P, Chen D, Collins M, Coulsen R, Davies R, Dyer PS, Farman M, Fedorova N, Feldblyum TV, Fischer R, Fosker N, Fraser A, Garcia JL, Garcia MJ, Goble A, Goldman GH, Gomi K, Griffith-Jones S, Gwilliam R, Haas B, Haas H, Harris D, Horiuchi H, Huang J, Humphray S, Jimenez J, Keller N, Khouri H, Kitamoto K, Kobayashi T, Konzack S, Kulkarni R, Kumagai T, Lafon A, Latge JP, Li W, Lord A, Lu C, Majoros WH, et al. 2005. Genomic sequence of the pathogenic and allergenic filamentous fungus *Aspergillus fumigatus*. Nature 438:1151-6.

3.	Fuchs BB, O'Brien E, El Khoury JB, Mylonakis E. 2010. Methods for using *Galleria mellonella* as a model host to study fungal pathogenesis. Virulence 1:475-482.

**Supplementary Figures**

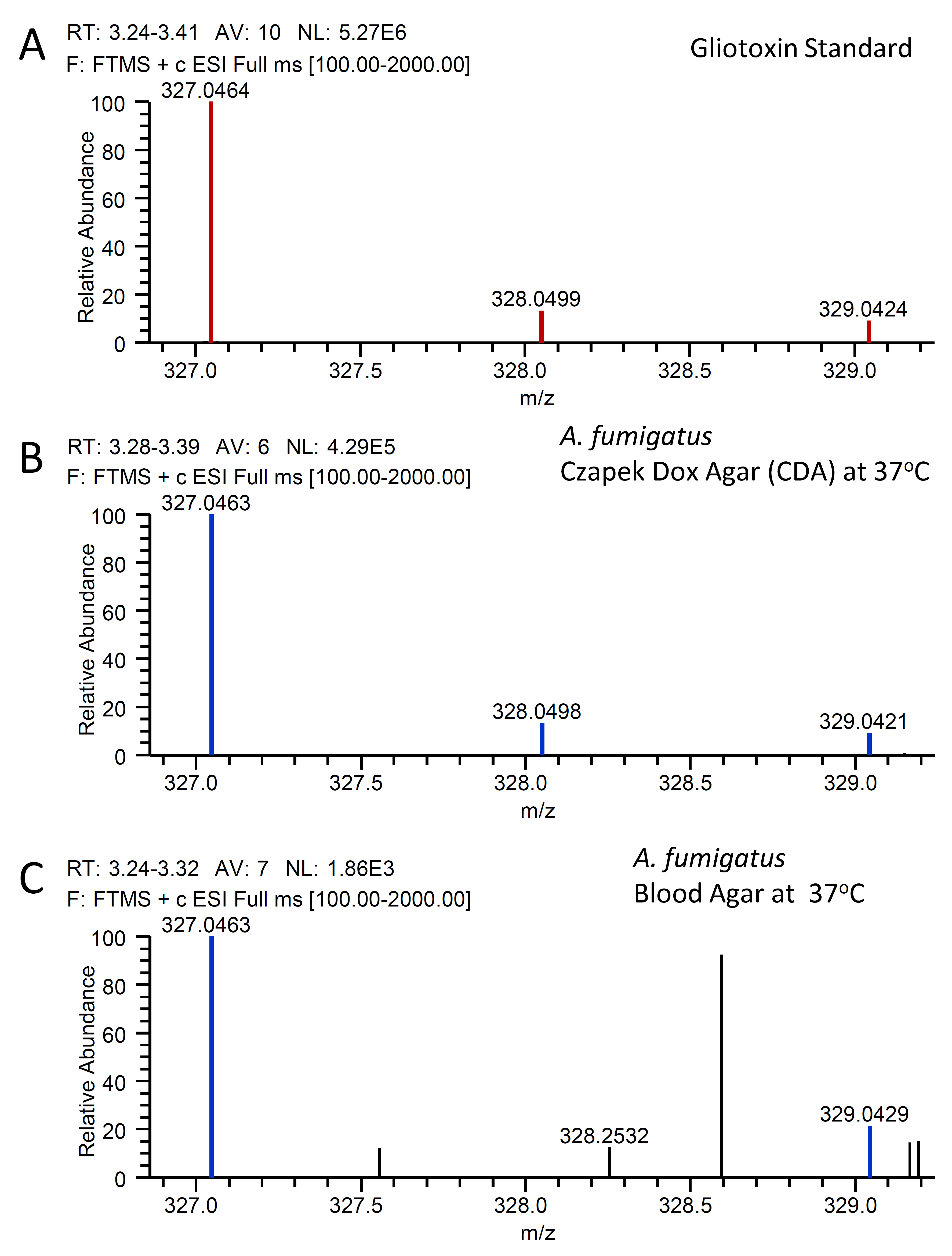

**FIG S1.** The mass spectra of gliotoxin in *A. fumigatus* grown on CDA and blood agar at 37°C verify the biosynthesis of gliotoxin by cultures of *A. fumigatus* on both CDA and blood agar at 37^o^C. Data are presented as mass to charge ratios (*m/z*). **A.** The isotopic pattern of the standard gliotoxin. **B.** The isotopic pattern of gliotoxin observed in *A. fumigatus* grown on CDA at 37^o^C. **C.** The isotopic pattern of gliotoxin observed in *A. fumigatus* grown on blood agar at 37^o^C.

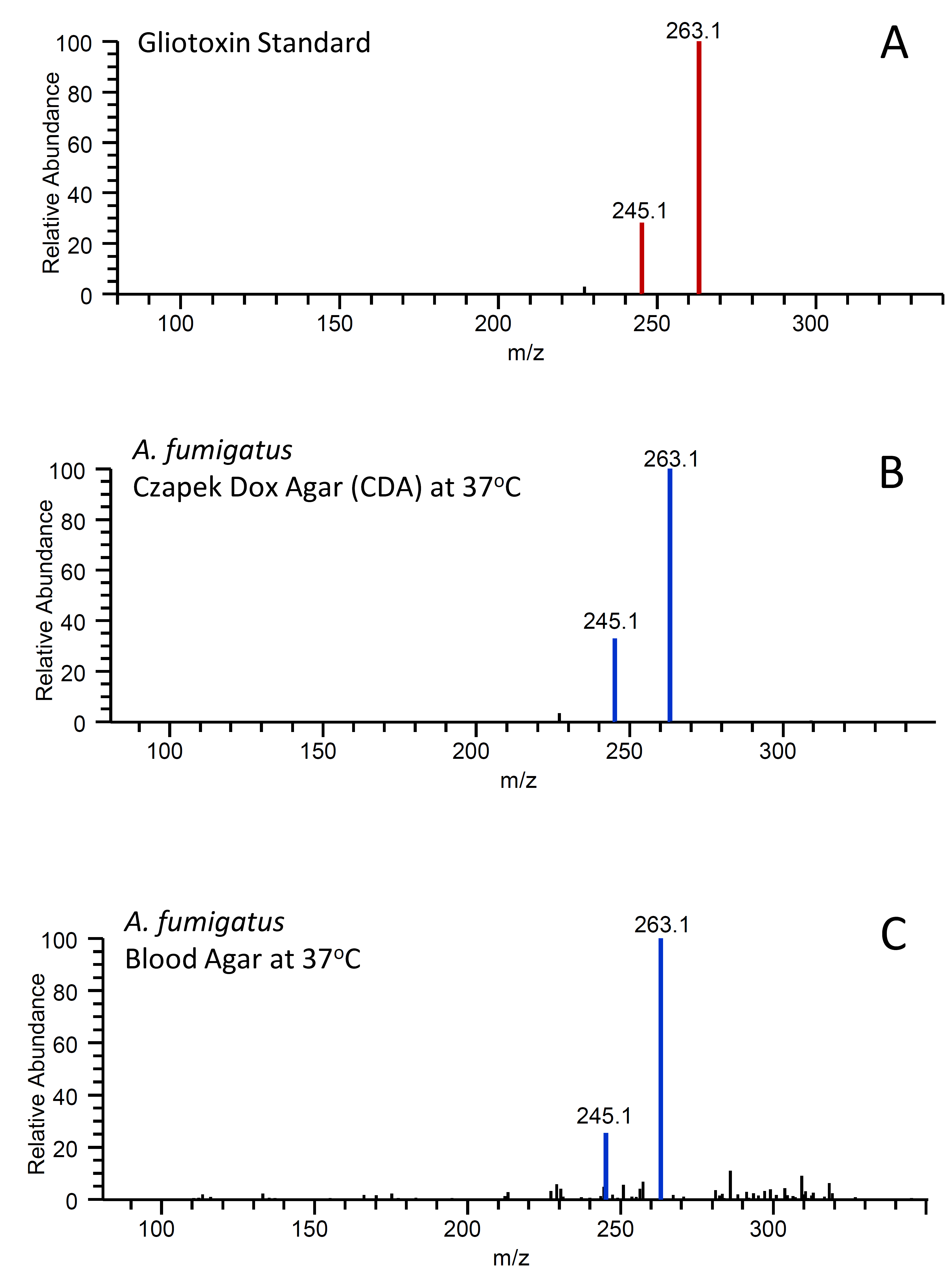

**FIG S2.** The fragmentation pattern (i.e., MS/MS data) of gliotoxin in *A. fumigatus* grown on CDA and blood agar at 37°C verify the biosynthesis of gliotoxin in both the CDA and blood agar growths of *A. fumigatus* at 37^o^C. **A.** The fragmentation pattern of the gliotoxin standard (263.1 and 245.1). **B.** The fragmentation pattern of gliotoxin observed in *A. fumigatus* grown on CDA incubated at 37^o^C. **C.** The fragmentation pattern of gliotoxin observed in *A. fumigatus* grown on blood agar incubated at 37^o^C.

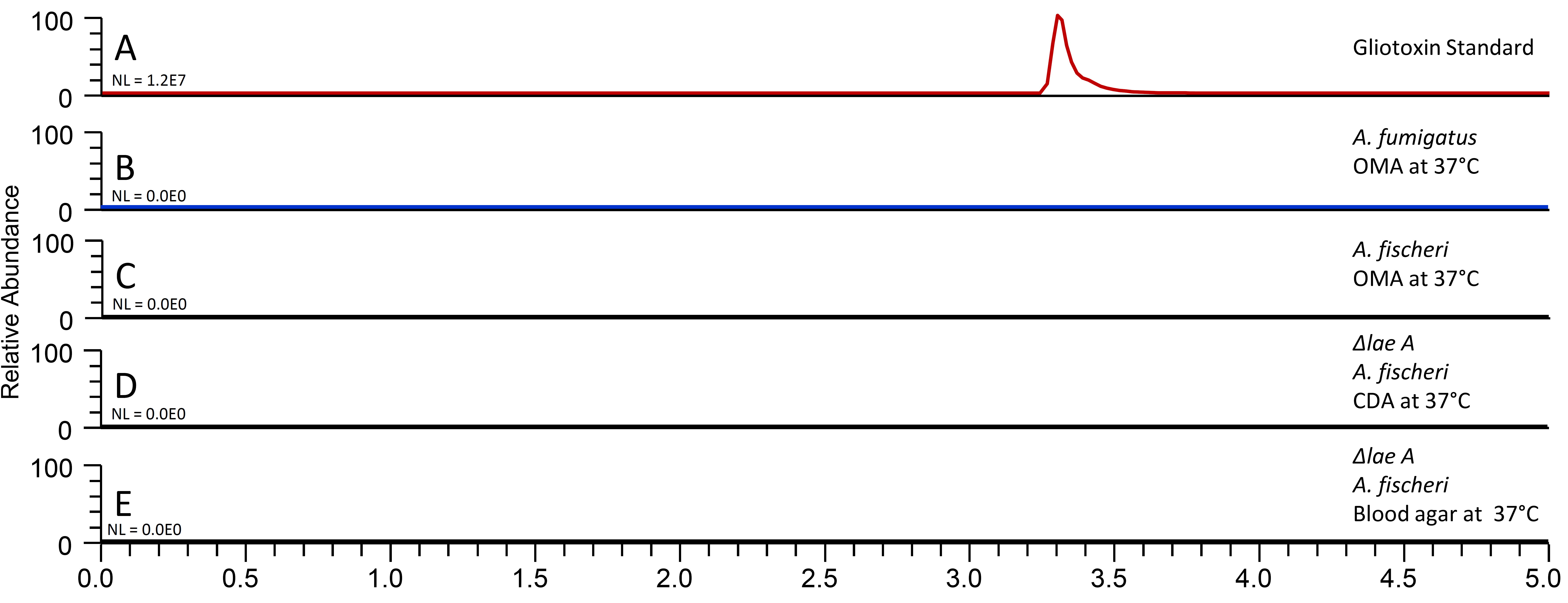

**FIG S3.** Base peak chromatograms of the gliotoxin standard, *A. fumigatus* grown on OMA at 37°C, *A. fischeri* grown on OMA at 37°C, *ΔlaeA A. fischeri* grown on CDA at 37°C, and *ΔlaeA A. fischeri* grown on blood agar at 37°C show that some media do not induce gliotoxin biosynthesis. Additionally, a negative control was analyzed (Panel C), to confirm that the observation of gliotoxin biosynthesis (**Fig. 1**) was genuine and not simply due to system carryover. Each sample (0.2 mg/mL) was analyzed by UHPLC-HRESIMS, and the data are presented as extracted ion chromatograms (XIC) using the protonated mass of gliotoxin (C_13_H_15_N_2_O_4_S_2_; [M+H]^+^ = 327.0473) and a window of ± 5.0 ppm. **A.** Analysis of the gliotoxin standard (0.01 mg/mL). **B.** *A. fumigatus* grown on OMA incubated at 37°C. **C.** *A. fischeri* grown on OMA incubated at 37°C. **D.** *ΔlaeA A. fischeri* grown on CDA incubated at 37°C. **E.** *ΔlaeA A. fischeri* grown on blood agar incubated at 37°C.

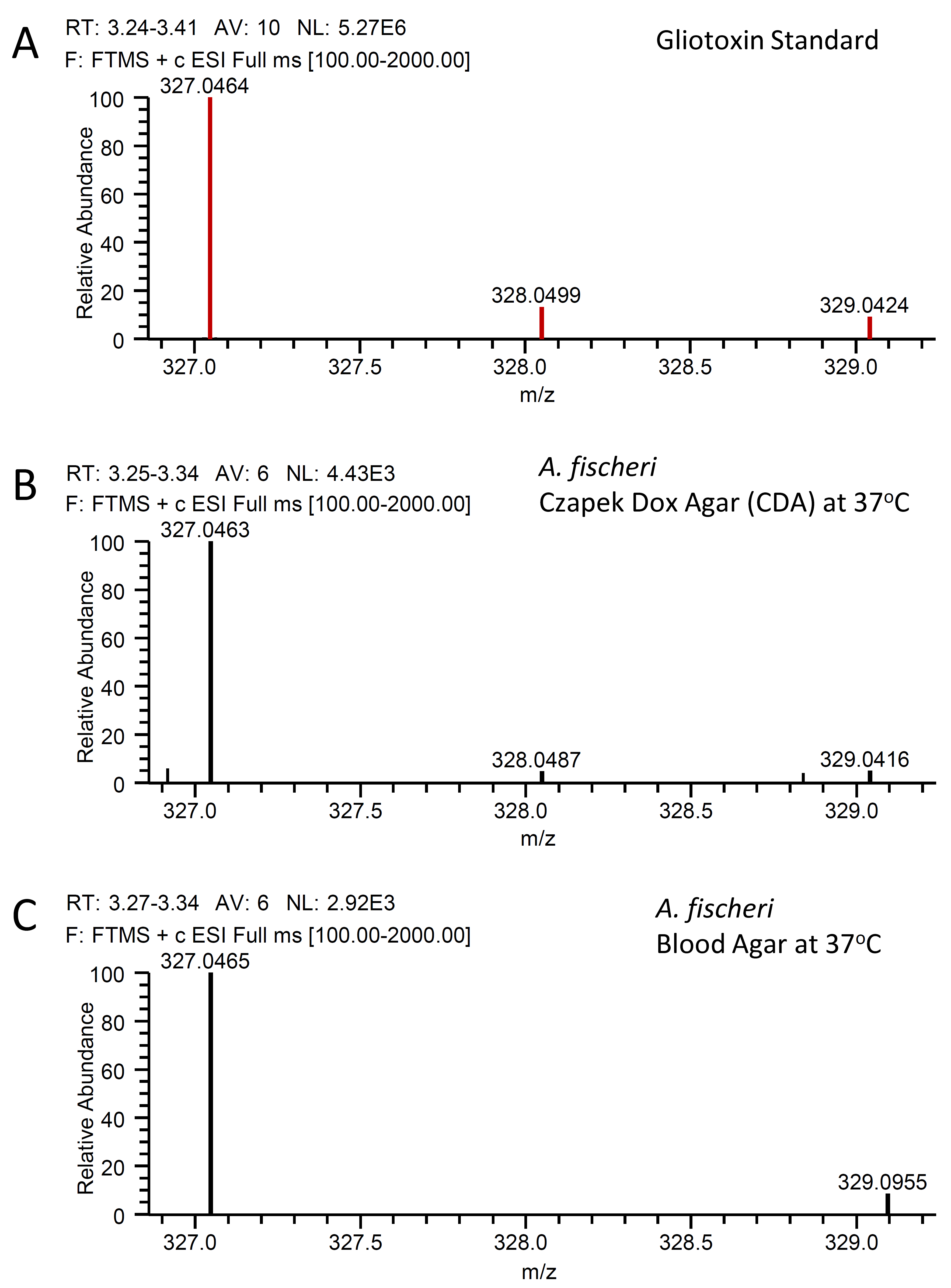

**FIG S4.** The mass spectra of gliotoxin in *A. fischeri* verify the biosynthesis of gliotoxin by cultures of *A. fischeri* on both CDA and blood agar at 37^o^C. The data are presented as mass to charge ratios (*m/z*). **A.** The isotopic pattern of the standard gliotoxin. **B.** Isotopic pattern of gliotoxin observed in *A. fischeri* grown on CDA at 37^o^C. **C.** Isotopic pattern of gliotoxin observed in *A. fischeri* grown on blood agar at 37^o^C.

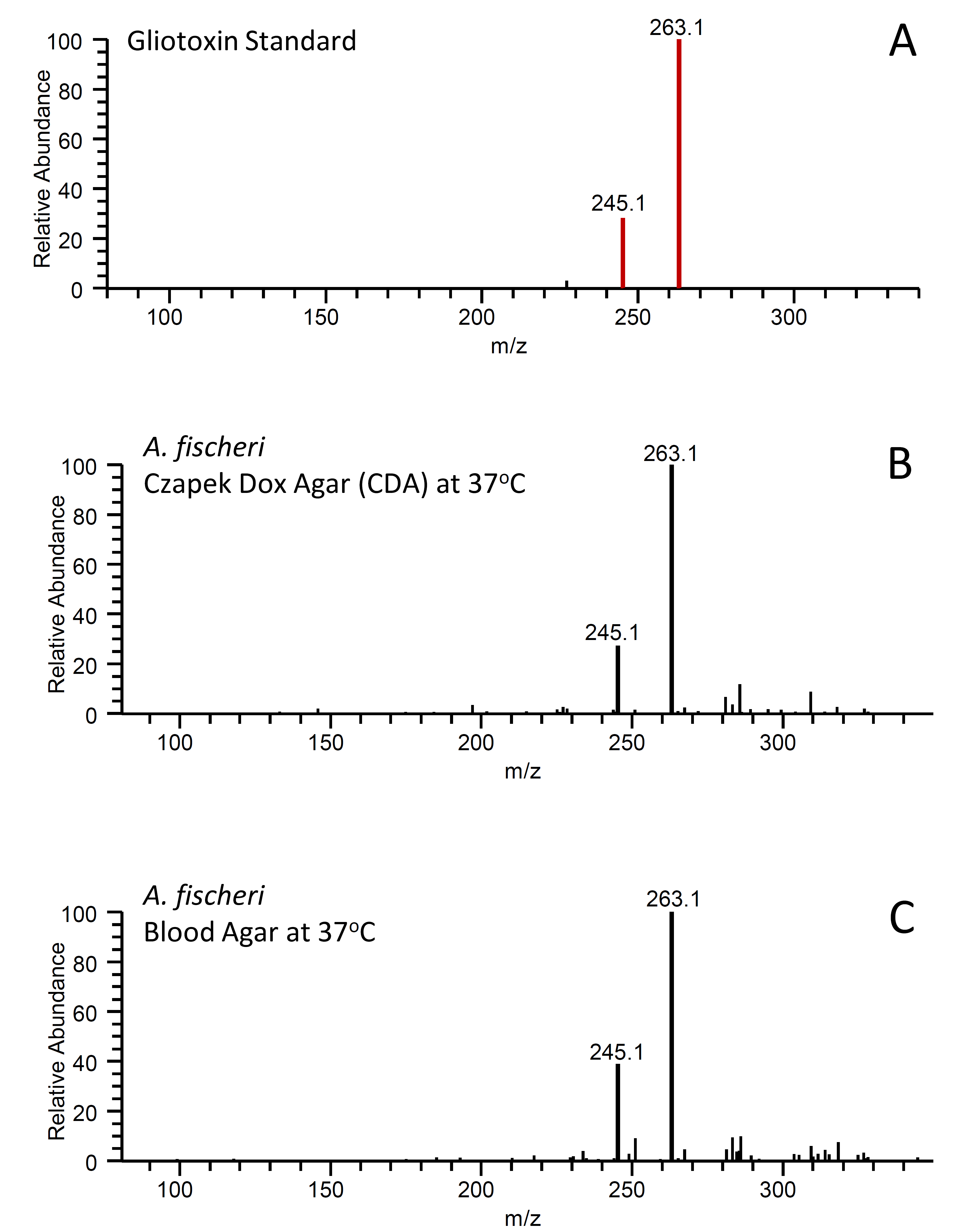

**FIG S5.** The fragmentation patterns (i.e., MS/MS data) of gliotoxin in *A. fischeri* further verify the biosynthesis of gliotoxin by cultures of *A. fischeri* in both the CDA and blood agar at 37^o^C. **A.** Fragmentation pattern of the gliotoxin standard (263.1 and 245.1). **B.** Fragmentation pattern of gliotoxin observed in *A. fischeri* grown on CDA incubated at 37^o^C. **C.** Fragmentation pattern of gliotoxin observed in *A. fischeri* grown on blood agar incubated at 37^o^C.
